# Integrative drug discovery for Parkinson’s disease with DIMER

**DOI:** 10.64898/2026.10.28.762666

**Authors:** Shuai Guo, Xingyi Yang, Yi Dong, Lidong Yu, Angela Sangjizhuoma, Ka Lok Sie, Keyun Zhu, Yiying Zhang, Xiyuan Zhao, Ella Kerman, Ruonan Li, Jing He

**Author notes:** Co-first authors (Shuai Guo, Xingyi Yang).

## Abstract

Translating Parkinson’s disease (PD) genetics into therapeutic candidates requires approaches that account for interconnected disease mechanisms and their cellular context. Here we present DIMER (**<u>D</u>**irect and **I**ndirect **<u>M</u>**odule **<u>E</u>**valuation for **<u>R</u>**epurposing and screening), a framework that connects network-based drug repurposing and target-directed virtual screening through shared disease-associated gene sets. Applying DIMER to PD, we integrated genetic risk and mechanism-associated gene sets with drug-target interactions, disease transcriptomes, and single nucleus expression profiles. Network analysis identified 423 repurposing candidates, with 71% of qualifying drug-gene set associations arising through targets outside the corresponding disease gene set. Transcriptional reversal analysis and cell type-resolved networks further distinguished candidates by their predicted effects on disease-associated expression and their cellular context, while target-set separation nominated potentially complementary drug combinations. For target-directed virtual screening, DIMER adapted a pretrained compound-protein interaction model to screening library chemistry and incorporated known-ligand similarity into compound ranking. Library adaptation improved compound-target pair classification performance, whereas ligand-informed ranking increased early enrichment of known active compounds. Consensus screening of 1.82 million compounds against seven PD-associated proteins yielded a shortlist of 22 compounds supported by at least five ranking runs. In a complementary experimental evaluation, four repurposing candidates (polydatin, fedratinib, sorafenib and venlafaxine) attenuated the loss of dopamine neuron marker signals in human midbrain organoids exposed to 6-hydroxydopamine. Together, these findings establish DIMER as a framework for translating disease-associated genes into pharmacological hypotheses and prioritizing candidates for experimental testing.

## Introduction

Parkinson’s disease (PD) is characterized by progressive degeneration of substantia nigra pars compacta (SNc) dopamine neurons and involves interconnected disturbances in protein homeostasis, mitochondrial function, oxidative stress, and inflammation^1–3^. Current therapies alleviate symptoms, but no approved treatment has been shown to slow disease progression^2^. Genetic and molecular profiling studies have identified disease-associated genes, pathways, and vulnerable cell populations^4,5^, creating opportunities to develop interventions grounded in PD biology. Translating these findings into therapeutic candidates, however, requires more than identifying individual disease genes. Many implicated proteins lack established pharmacological modulators, and the effects of targeting them depend on their molecular interactions and cellular context. A central challenge is therefore to connect disease-associated biology to compounds that can influence it, whether through existing drugs acting on connected proteins or new compounds directed at selected disease-associated targets.

Network medicine offers a framework for making these connections by placing disease-associated genes within the human protein-protein interaction (PPI) network^4,6–8^. Network proximity identifies drugs whose targets lie near a disease module, including through interacting proteins^8^. This approach has nominated repurposing candidates for neurodegenerative diseases^4,9^ and aging^6^. Network relationships can also inform combination design: target-set separation identifies drug pairs that approach the same disease module through distinct network neighborhoods^7^, providing a rationale for investigating complementary interventions. However, network proximity establishes a connection to disease biology without specifying its therapeutic direction^6^ or relevance to affected cell types^10,11^. A proximal drug may oppose or reinforce disease-associated molecular changes^6^, and its relevant targets or interactions may be restricted to particular cell populations^10,11^. Integrating drug-induced transcriptional responses^12^ with cell type-resolved expression data can therefore help distinguish how, and in which cellular settings, a candidate may influence a disease module.

AI-based virtual screening complements network-based drug repurposing by expanding the search beyond drugs with known targets. Disease-associated networks can guide target selection by identifying highly connected proteins whose genetic, molecular, and pharmacological evidence supports further investigation^13^. Models, such as AI-Bind^14^, ColdstartCPI^15^, DrugBAN^16^, and DrugCLIP^17^, predict compound-protein interactions from molecular structures and protein sequences/structures, enabling large chemical libraries to be screened without requiring experimentally established interactions for each candidate. Applying these models to specialized libraries, however, introduces additional challenges. Libraries selected for physicochemical properties favorable for central nervous system exposure^18,19^ may occupy a different chemical space from the model’s training data, motivating adaptation to the intended screening library. Moreover, accurate compound-protein pair classification does not necessarily ensure that active compounds rank highly for an individual target^20^. Where experimentally established ligands are available, their chemical similarity to screened compounds can provide additional target-specific information for prioritization^21^. Combining disease-informed target selection, library adaptation, and ligand-informed ranking therefore offers a strategy for identifying new chemical candidates against proteins implicated in PD.

Another challenge for both approaches is the experimental validation of computationally nominated compounds in a human-relevant system. Cell lines such as SH-SY5Y provide a scalable platform for rapid screening but do not capture the cellular diversity or three-dimensional organization of the human midbrain^22^. Animal models, including α-synuclein preformed fibril-inoculated^23^ and neurotoxin-induced models using MPTP^24^ or 6-OHDA^25^ offer anatomical and physiological complexity but are costly and time-consuming. Human midbrain organoids preserve key aspects of tissue architecture and cellular diversity and support genetic, protein aggregation, and toxin-induced PD models^26–28^. They therefore provide a practical, human-relevant platform for evaluating whether computationally prioritized compounds exert neuroprotective effects.

Here we introduce DIMER (**<u>D</u>**irect and **I**ndirect **<u>M</u>**odule **<u>E</u>**valuation for **<u>R</u>**epurposing and screening), a framework that connects network-based drug repurposing and target-directed virtual screening through a shared representation of disease biology. We applied DIMER to PD using 175 genetic risk genes and complementary 508 disease-and mechanism-associated gene sets from Open Targets. We separated the 508 genes from Open Targets into six modules representing PD-related biological processes. For repurposing, DIMER integrates network proximity with transcriptional reversal, assessed using RevERT (**Rev**ersal of **E**xpression **R**elative to the disease **T**ranscriptome), and cell type-resolved networks to characterize candidate drugs by their disease connections, predicted transcriptional effects, and cellular context. Target-set separation additionally nominates potentially complementary drug partners. For virtual screening, we adapt the pretrained AI-Bind model to the chemical properties of the screening library, and incorporate similarity to known ligands into compound ranking. Benchmarks showed improvements in compound-protein pair prediction and enrichment of known active compounds. We then screened 1.82 million compounds against seven prioritized PD-associated proteins using DMIER and five existing predictors: AI-Bind^14^, ColdstartCPI^15^, DrugBAN^16^, DrugCLIP^17^, and GraphDTA^29^. We evaluated four selected repurposing drugs in human midbrain organoids exposed to 6-hydroxydopamine (6-OHDA). By using disease-associated gene sets as a shared input, DIMER connects these complementary network-proximity and virtual-screening strategies within a general framework that could be potentially applied to other diseases.

## Results

### DIMER targets a disease module indirectly and directly

We developed DIMER (**<u>D</u>**irect and **I**ndirect **<u>M</u>**odule **<u>E</u>**valuation for **<u>R</u>**epurposing and screening), an integrative framework that uses disease-associated gene modules to prioritize therapeutic candidates from annotated drugs and chemical libraries (Fig. 1a, see Methods). DIMER has two complementary strategies. The indirect strategy combines a curated human PPI network^4,6–8^ with drug-target interaction (DTI) annotations to identify drugs whose targets lie close to disease genes. The PPI network^6^ used for analysis contains 18,208 proteins and 524,155 interactions, and our integrated DTI library^30–36^ comprises 2,651 drugs to 3,810 target proteins. DIMER uses network proximity to identify drugs whose targets lie close to disease genes. It then uses a metric named RevERT (**Rev**ersal of **E**xpression **R**elative to the disease **T**ranscriptome) to compare drug perturbation signatures with disease-associated expression changes and estimate the direction of each drug’s transcriptional effect^6^. Target-set separation identifies partners for existing therapies^7^, with documented drug-drug interactions^7,37^ considered during prioritization. Single cell data^5^ allow proximity and RevERT to be evaluated across cell types. The direct strategy selects druggable hub proteins from the disease module and screens chemical libraries using AI-Bind^14^ adapted to the library’s chemical properties. We inform compound ranking based on similarity to known ligands^21^ and choose consensus ranking candidates supported by multiple prediction models^14–17,29^. Candidates prioritized by DIMER can be evaluated in human organoid models^38^. In this study, we tested selected repurposing drugs in human midbrain organoids exposed to 6-hydroxydopamine^26,27^ (6-OHDA).

**Fig. 1.**
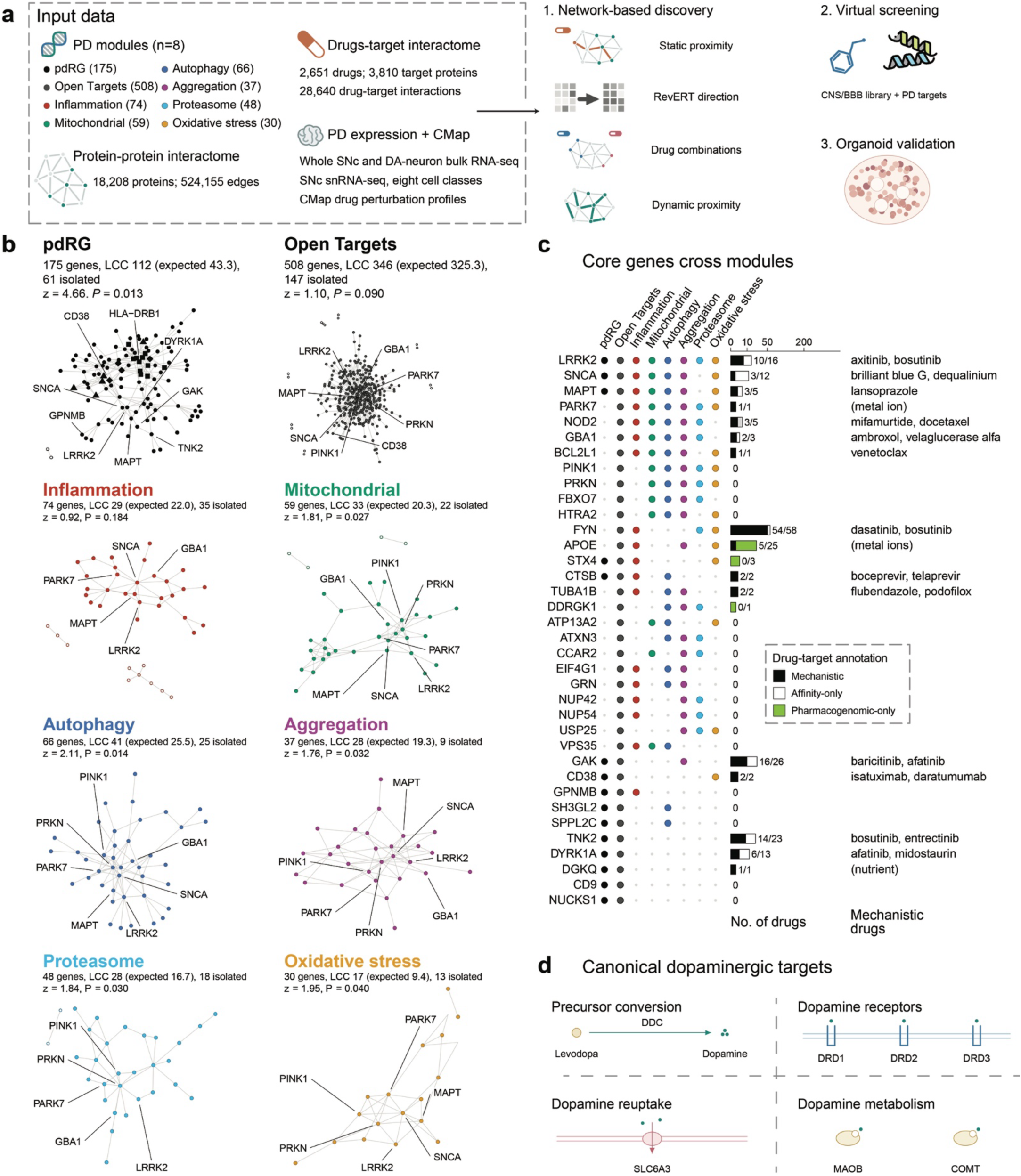
DIMER framework and the PD gene modules. **a,** Overview of DIMER framework. **b,** PPI subgraphs of the pdRGs, Open Targets, and six PD-associated gene sets. Filled nodes belong to the LCC; open nodes lie outside it. Headers report set size, observed and expected LCC size, isolated genes, z and one-sided empirical P from 1,000 degree-matched random sets. **c,** Membership and drug-target coverage of 36 shared core genes. Bars show drug-target annotations; numbers indicate drugs with mechanistic evidence out of all drugs annotated to each gene. **d,** Canonical dopaminergic processes and targets of symptomatic therapy. PD, Parkinson’s disease; CMap, Connectivity Map; SN, substantia nigra; DA, dopaminergic; SNc, substantia nigra pars compacta; CNS, central nervous system; BBB, blood-brain barrier.

We applied DIMER to PD using eight gene sets representing genetic evidence and PD-related biological processes (Fig. 1b). These included 175 PD risk genes (pdRGs) prioritized from genome-wide association and functional-genomic evidence^4^, 508 genes with genetic associations to PD from Open Targets^39^, and six mechanism sets of 30-74 genes derived from the Open Targets genes using pathway annotations^1^: inflammation, mitochondrial dysfunction, autophagy, protein aggregation, proteasome dysfunction, and oxidative stress. To assess their network connectivity, we compared the largest connected component (LCC) of each set’s interactome subgraph with that of degree-matched random gene sets. The pdRGs formed an LCC of 112 genes, 2.6-fold of the 43.3 genes expected by chance (z = 4.66, *P* = 0.013; Fig. 1b). Autophagy also showed LCC enrichment (z = 2.11, *P* = 0.014), with more modest enrichment for mitochondrial dysfunction, protein aggregation, proteasome dysfunction, and oxidative stress (z = 1.76-1.95, *P* = 0.027-0.040). Enrichment was not significant for inflammation (z = 0.92, *P* = 0.184) or the full Open Targets set (z = 1.10, *P* = 0.090). We retained all eight sets to represent different aspects of PD biology.

The gene sets shared a core of 36 genes, defined as genes present in at least three mechanism sets or in both major genetic sets (i.e., 175 pdRGs and 508 Open Targets; Fig. 1c). *LRRK2* belonged to all eight PD-associated sets, followed by *SNCA*, *MAPT*, and *PARK7* who belonged to seven gene modules. Half of the core genes (18 of 36) had no direct drug-target annotation in our curated DTI interactome, including well-known PD-associated genes such as *PINK1*, *PRKN*, and *VPS35*. Among genes with drug-target annotations, *LRRK2* was linked to kinase inhibitors such as axitinib and bosutinib, and *GBA1* to ambroxol and velaglucerase alfa.

The targets of established symptomatic PD therapies, including DRD1-DRD3, DDC, SLC6A3, MAOB and COMT, were absent from all eight gene sets (Fig. 1d). These targets are associated with drugs such as levodopa, pramipexole, carbidopa, selegiline, and entacapone. Together, we show partial annotated drug-target coverage of the shared PD gene core and support evaluating drugs whose targets connect to disease genes through the PPI network.

### Static network proximity nominates repurposing drug candidates

We calculated static network proximity between the targets of 2,651 drugs in our DTI library and each of the eight PD gene sets. After applying drug eligibility and target-evidence criteria (see Methods), we identified 423 candidates proximal to at least one gene set, comprising 766 drug-module pairs (z ≤ -1.96 and empirical *P* < 0.05; Fig. 2a,b). The four largest groups were kinase inhibitors, adrenergic agents, antibody or protein biologics, and monoamine reuptake inhibitors or releasers, together accounting for 177 candidates. Among the 423 proximal candidates, 247 were proximal to only one PD-associated gene set, and 176 were proximal to 2-6 gene sets (Fig. 2a).

**Fig. 2.**
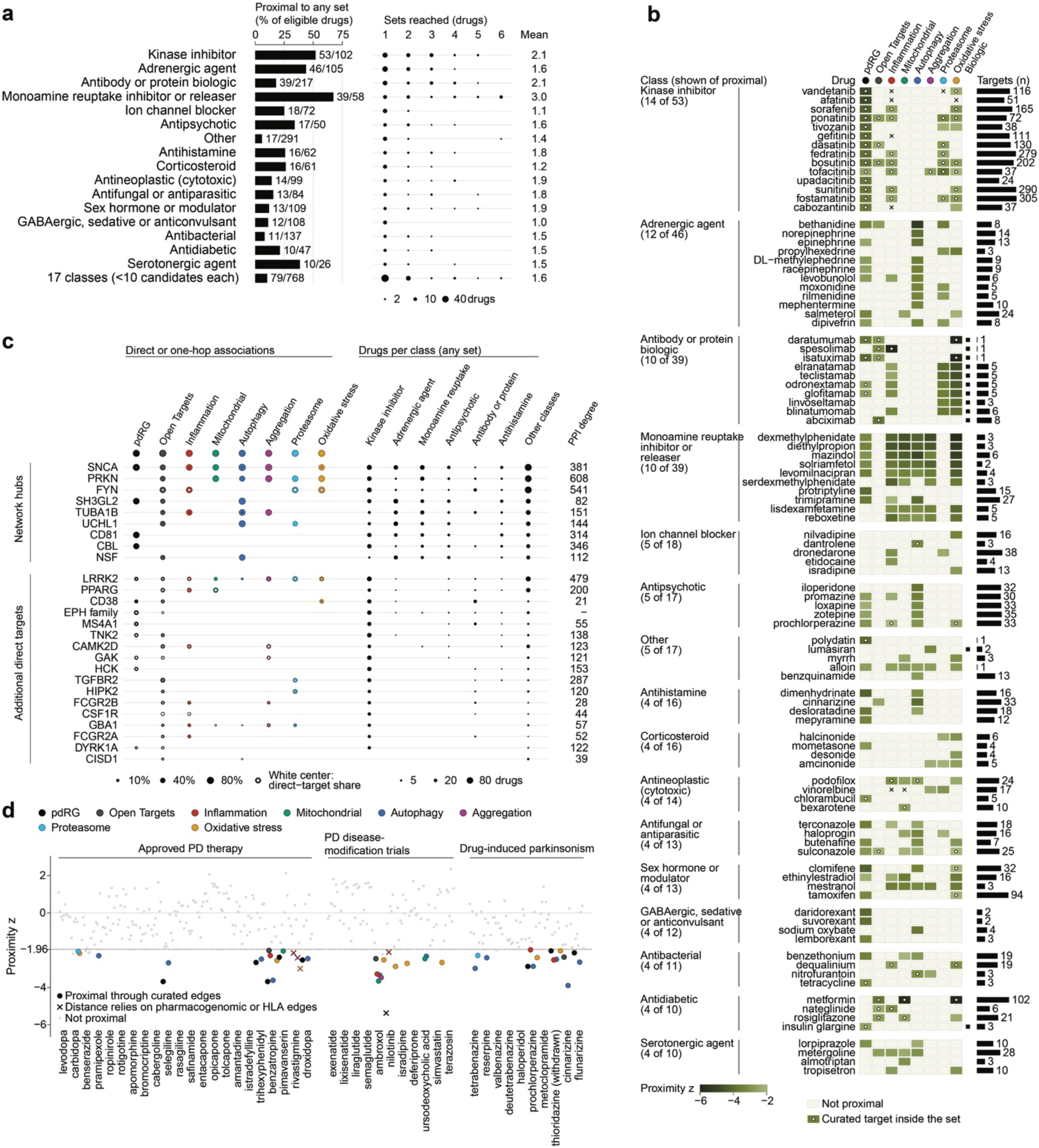
Static network associations of drugs with PD modules. **a,** Percentage of eligible drugs proximal to at least one PD gene set, and number of gene sets reached, by drug class. **b,** Representative drug-module associations. Darker color indicates stronger proximity; white dots mark targets within the module; crosses indicate dependence on pharmacogenomic-only or HLA targets; black squares identify biological drugs, i.e., antibodies and therapeutic proteins. Bars show target counts. **c,** Module genes connected to drugs directly or through one PPI interaction with a target outside the module. Dot size shows the fraction of each module’s proximal candidates connected to the gene; white centers show the directly targeting fraction. Black dots show drug counts by class. **d,** Profiles of 22 approved PD therapies, 11 disease-modification trial drugs and ten drugs associated with parkinsonism.

Candidates proximal to the pdRGs set included kinase inhibitors targeting EPH receptors, GAK, LRRK2, and DYRK1A, and antibodies targeting CD38 or MS4A1 (Fig. 2b,c). Examples included sorafenib (z = -4.21, empirical *P* = 0.014) and fedratinib (z = -3.84, empirical *P* = 0.014). Polydatin was also proximal to the pdRG set (z = -3.99, empirical *P* = 0.017). The mechanism sets identified additional associations (Fig. 2b). CSF1R and JAK inhibitors were proximal to the inflammation set. Metformin was proximal to mitochondrial dysfunction (z = -5.78, empirical *P* = 0.004), the adrenergic agent bethanidine to autophagy (z = -5.28, empirical *P* = 0.001), and the anti-CD38 antibody daratumumab to oxidative stress (z = -6.90, empirical P = 0.002). Fedratinib was proximal to inflammation and proteasome dysfunction, and sorafenib to inflammation and oxidative stress, showing that one candidate could be associated with several PD modules.

About 71% of prioritized drug-module pairs (546 of 766) involved drugs whose curated targets were outside the corresponding PD gene set. These drugs were identified through indirect connections to the modules, as their targets were close to module genes in the PPI network. The most frequently connected module genes included *SNCA*, *PRKN*, *FYN*, and *SH3GL2* (Fig. 2c). For example, monoamine transporter-targeting drugs, such as desvenlafaxine, levomilnacipran, and dexmethylphenidate, act on transporter proteins (SLC6A2, SLC6A3, and SLC6A4) that interact with *SNCA* and *PRKN* in the network.

We next examined the proximity profiles of three reference groups^40,41^: established PD therapies (n = 22), drugs evaluated in PD disease-modification trials (n = 11), and drugs associated with drug-induced parkinsonism (n = 10) (Fig. 2d). Seven of 22 established PD therapies, five of 11 disease-modification trial drugs, and six of 10 parkinsonism-associated drugs were proximal to at least one PD module. Ambroxol was proximal to six gene sets. Tetrabenazine and reserpine, both associated with drug-induced parkinsonism^42^, were proximal to the autophagy set. These results show that drugs used to modify PD symptoms and drugs that can cause parkinsonism can both be proximal to PD-associated gene sets.

### RevERT evaluates the direction of drug-induced transcriptional changes

Network proximity alone does not establish whether a drug’s effects are beneficial or harmful. We therefore developed RevERT to provide an estimate of transcriptional direction by assessing whether drug-induced expression changes oppose or reinforce disease-associated changes. For each PD module, RevERT compares Connectivity Map (CMap)^43^ drug perturbation signatures with disease-vs.-control expression changes among measured module genes. Genes with stronger statistical evidence of disease-associated changes receive greater weight, and scores are averaged across each drug’s CMap signatures (see Methods). Positive scores indicate reversal, and negative scores indicate reinforcement. We calculated RevERT for 1,344 drugs using disease-expression references from whole substantia nigra (SNc)^44^ and laser-captured dopamine (DA) neurons^45^.

The two references showed a weak correlation in differential-expression t statistics across 14,311 shared genes (Pearson *r* = 0.14; Fig. 3a). *TH* expression was lower in PD than controls in whole SNc^44^ but showed little change in the sampled dopamine neurons^45^. In these neurons, *GPNMB* and *HSP90AA1* expression was higher in PD than in controls. Among 55 module landmark genes measured in both references, 25 showed opposite directions of expression change (Fig. 3b).

**Fig. 3.**
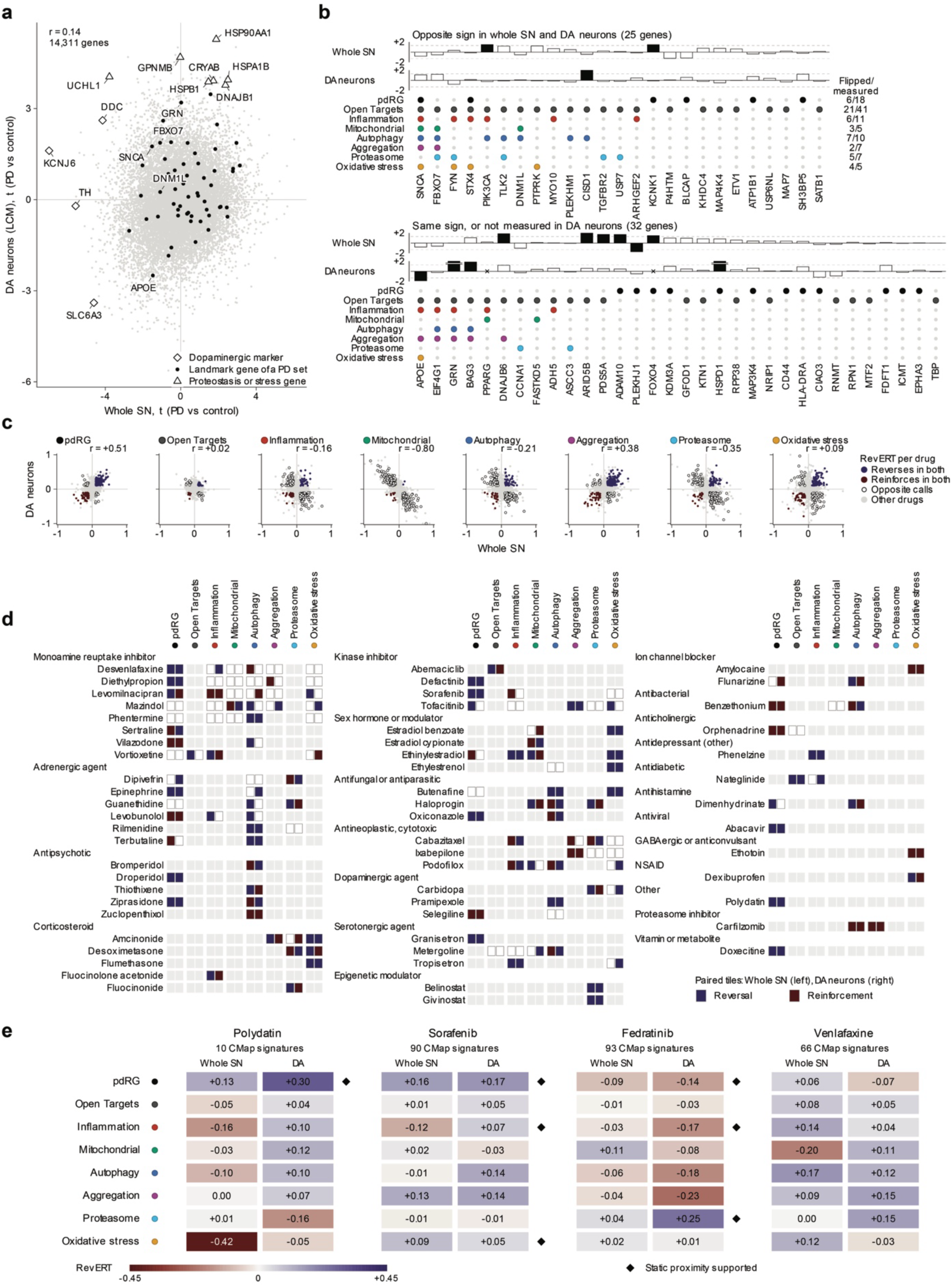
RevERT profiles across whole tissue and dopamine neuron disease references. **a,** Scatter plot shows PD-vs-control t statistics for 14,311 shared genes in whole SN and laser-captured DA neurons. **b,** Dot plots shoes disease-direction weights, sign (log_2_ fold change) × -log_10_(*P*), and module membership of 57 CMap landmark genes. **c,** Scatter plots show Cross-reference RevERT scores for 1,344 drugs per module. Positive scores indicate reversal and negative scores reinforcement. Blue and red points meet the corresponding direction criteria in both references; open circles have opposite calls. **d,** Profiles of 59 drugs with a direction call in both references for at least one eligible proximal pair. Paired tiles show whole SN and DA: blue, reversal; red, reinforcement; white, no call; grey, outside the eligible proximal set. **e,** Continuous RevERT profiles of the four organoid-tested compounds; diamonds mark supported static proximity. Direction calls require at least three signatures, |mean RevERT| ≥ 0.1 and ≥ 60% sign agreement.

RevERT scores calculated using the two references were positively correlated for pdRGs (Pearson *r* = 0.51) and protein aggregation (Pearson *r* = 0.38) and negatively correlated for mitochondrial dysfunction (Pearson *r* = -0.80; Fig. 3c). In the mitochondrial set, *SNCA*, *FBXO7*, and *DNM1L* changed in opposite directions between references and accounted for most of the gene weight used to calculate RevERT. Of 495 eligible proximal drug-module pairs with RevERT scores, 30 met the reversal criteria in both references (Fig. 3d). Examples included tofacitinib for protein aggregation, givinostat for proteasome dysfunction, and pramipexole for autophagy. Their mean RevERT scores were +0.22/+0.26, +0.30/+0.69, and +0.12/+0.18 in whole SN/DA neurons, respectively. Carfilzomib reinforced the protein aggregation-associated expression changes in both references, with scores of -0.24 and -0.25.

The four compounds subsequently tested in organoids showed various RevERT profiles (Fig. 3e). Polydatin and sorafenib met the reversal criteria for pdRGs in both references, with scores of +0.13/+0.30 and +0.16/+0.17, respectively. Using the dopamine neuron reference^45^, fedratinib showed reversal for proteasome dysfunction and reinforcement for pdRGs and inflammation. Venlafaxine met the reversal criteria for autophagy in both references (+0.17/+0.12), although it did not meet the static proximity thresholds.

### Complementary drug pairs reach PD modules through separated target neighborhoods

To identify drugs that could complement established PD therapies, we combined module proximity with target-set separation^7,37^. We sought pairs whose targets occupied distinct network neighborhoods close to the same disease module. Across 2,651 drugs, we evaluated all 3,512,575 pairs. Among pairs in which both drugs were proximal to pdRGs, 24,423 had separated target neighborhoods (s ≥ 0), and 6,205 had overlapping target neighborhoods (s < 0; Fig. 4a).

**Fig. 4.**
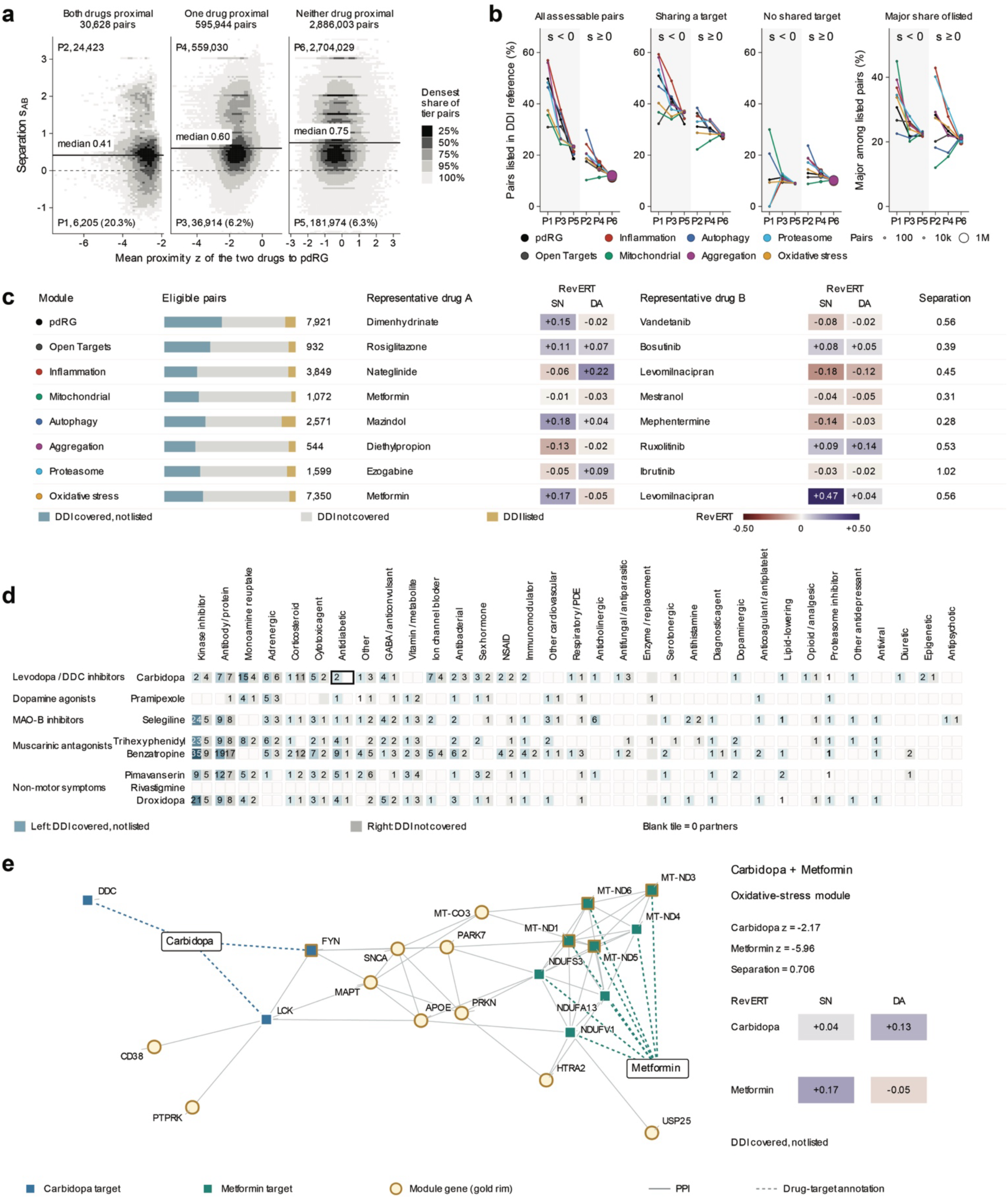
Network complementarity and potential partners for established PD therapies. **a,** Target-set separation among drug pairs with both, one, or neither drug proximal to pdRG. Greyscale regions contain 25%, 50%, 75%, 95% and 100% of pairs within each proximity tier; solid lines mark medians and the dashed line zero separation. Proximity requires z ≤ -1.96 and empirical *P* < 0.05 from 1,000 permutations. **b,** Documented DDIs across proximity and separation groups, with comparisons by shared targets and interaction severity. **c,** Candidate pairs across PD modules, with DDI annotations, representative combinations, and RevERT scores for each drug. **d,** Candidate partners for established PD therapies, grouped by drug class and DDI status. Pairs with listed DDIs are excluded. **e,** Carbidopa and metformin connect to the oxidative-stress module through separated target neighborhoods.

We next examined whether these network patterns were associated with documented drug-drug interactions (DDIs). We compared drug pairs where both drugs were proximal to the pdRGs and represented in the DDI reference. Adverse interactions were listed for 49.8% of overlapping pairs and 18.7% of separated pairs. This difference was much smaller among pairs without shared targets, with corresponding proportions of 10.4% and 13.0% (Fig. 4b).

We selected drug pairs to be proximal to the same module, curated target support, positive target-set separation (see Methods), and no shared targets, yielding 22,551 distinct candidate pairs across the eight gene sets. Examples included dimenhydrinate with vandetanib for pdRGs, and metformin with levomilnacipran for oxidative stress (Fig. 4c). Individual RevERT scores further described the transcriptional direction of each drug (Fig. 4c, left panel). For example, metformin and levomilnacipran both met the reversal criteria for the whole SN reference, with RevERT scores of 0.17 and 0.47, respectively.

We then examined candidate partners for established PD therapies by drug class (Fig. 4d). For carbidopa, which is frequently used with levodopa^46^, candidate partners included the antidiabetic drugs metformin and rosiglitazone. Carbidopa and metformin were both proximal to the oxidative-stress module (z = -2.17 and -5.96, respectively), with no shared targets and positive target-set separation (s = 0.706; Fig. 4e). Their network connections to the module involved annotated *FYN* and *LCK* targets for carbidopa, and mitochondrial complex I targets for metformin. Both drugs were represented in the DDI reference, and their combination was not listed as an adverse interaction.

### Context-specific networks refine candidates across substantia nigra

To assess whether drug-disease associations differed across cell types, we incorporated a single nucleus atlas^5^ of 290,017 SNc nuclei from seven PD and eight control donors into the proximity analysis (Fig. 5a). PPI connections were retained when both genes were detected in at least 10% of nuclei within a cell type, and edge lengths were adjusted using gene detection fractions. Drug targets and disease genes were weighted by cell type specificity and the magnitude of PD-associated expression changes (see Methods).

**Fig. 5.**
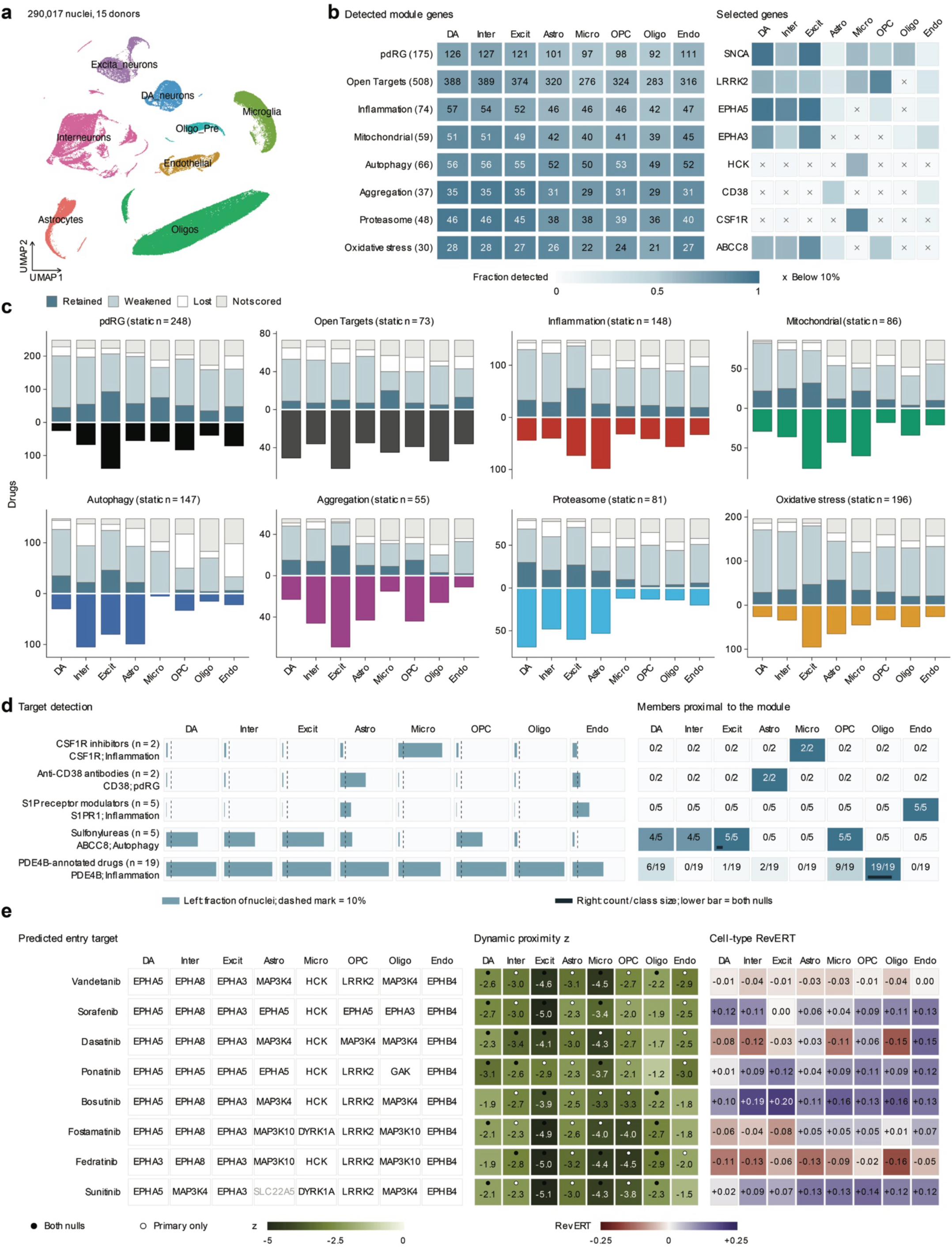
Cell-context-resolved proximity, target entry points and transcriptional direction. **a,** UMAP of 290,017 SNpc nuclei from PD (n=7) and control (n=8) donors, colored by cell classes. **b,** Module-gene detection and selected-gene expression; the detection threshold is 10% of nuclei. **c,** Changes in static drug candidates after cell type-specific proximity analysis. **d,** Target detection and qualifying drug-class fractions; dark lower bars indicate support under both nulls. **e,** Predicted entry targets, proximity, and RevERT for eight kinase inhibitors against pdRGs across cell types.

Gene detection varied across the eight cell classes. For example, 126 of the 175 pdRGs passed the detection threshold in dopamine neurons, compared with 92 in oligodendrocytes (Fig. 5b). Incorporating this cellular context changed the candidate sets. Of 248 drugs meeting the static proximity thresholds for pdRG, 45 remained proximal in dopamine neurons. In excitatory neurons, 93 remained proximal, and 139 additional drugs met the proximity thresholds (Fig. 5c).

Drug classes showed different proximity patterns across cell types (Fig. 5d). Both CSF1R inhibitors were proximal to inflammation in microglia, both anti-CD38 antibodies were proximal to pdRG in astrocytes, and all five sphingosine-1-phosphate (S1P) receptor modulators were proximal to inflammation in endothelial cells. Sulfonylureas were proximal to autophagy in neuronal populations and oligodendrocyte precursor cells, consistent with ABCC8 detection in these populations. All 19 PDE4B-annotated drugs were proximal to inflammation in oligodendrocytes, although PDE4B was broadly detected across cell classes. Fifteen of these associations also met the proximity criteria when the null model additionally matched PD-associated expression changes.

Kinase inhibitors connected to pdRGs through different annotated targets across cell types (Fig. 5e). Vandetanib was proximal to pdRGs in all eight cell classes. Its nearest targets to the module included EPHA5 in dopamine neurons, HCK in microglia, and LRRK2 in oligodendrocyte precursor cells. We also calculated cell type RevERT by comparing the same CMap drug signatures with the PD-versus-control expression signature of each cell type. Sorafenib was proximal to pdRGs in both dopamine neurons and excitatory neurons (z = -2.66 and -5.02, respectively). Its RevERT score was positive in dopamine neurons (+0.12) but close to zero in excitatory neurons, indicating different transcriptional directions despite proximity in both cell types.

### Library adaptation and ligand-informed ranking improve distinct screening tasks

For virtual screening, we adapted AI-Bind^14^ to the chemical properties of the screening library using two approaches (see Methods). Property-box adaptation fine-tuned the model on training compounds within the library’s physicochemical range. Density-ratio adaptation reweighted training compounds to match the library’s chemical distribution. Both approaches used the library’s chemical properties without requiring binding labels for its compounds. We also introduced similarity-band ranking, which orders compounds by similarity to known ligands and uses AI-Bind predictions to rank compounds within each similarity band (see Methods). We evaluated compound-protein pair prediction and enrichment of known active compounds in target-specific screening on DIMER and existing five predictors (AI-Bind^14^, ColdstartCPI^15^, DrugBAN^16^, DrugCLIP^17^, and GraphDTA^29^; Fig. 6a).

**Fig. 6.**
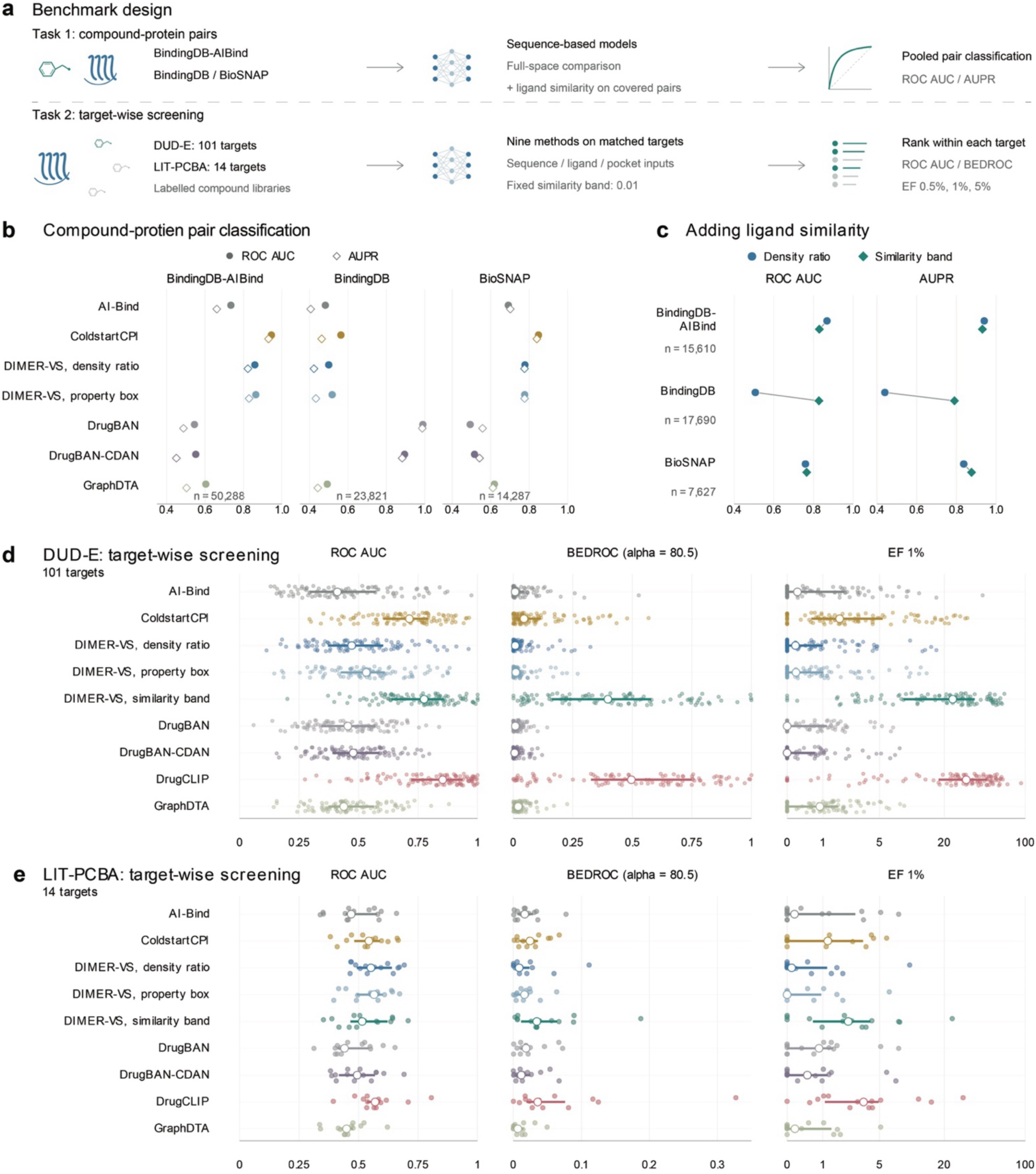
Benchmarking sequence-based adaptation and ligand-similarity integration. **a,** Evaluation of compound-protein pair prediction and enrichment of active compounds in target-specific screening. **b,** Pair-prediction performance of six models across three datasets. **c,** Contribution of ligand similarity, evaluated on the same pairs for proteins with at least ten reference ligands. **d and e,** Ranking and enrichment of known active compounds for 101 DUD-E targets **(d)** and 14 LIT-PCBA targets| **(e)**. Summaries show medians and interquartile ranges across targets.

Both adaptation approaches improved ROC AUC and AUPR on three benchmark datasets: BindingDB-AI-Bind^14^, BindingDB^34^, and BioSNAP^47^ (Fig. 6b). On BindingDB-AIBind and BioSNAP, DIMER property-box and density-ratio adaptation increased ROC AUC from 0.734 to 0.864 and 0.859, respectively, and AUPR from 0.661 to 0.829 and 0.823. Improvements were smaller on BindingDB.

We next assessed the contribution of ligand similarity using proteins with at least ten known reference ligands. Density-ratio adaptation and similarity-band ranking were compared on the same compound-protein pairs within this subset (Fig. 6c). On BindingDB, incorporating ligand similarity increased ROC AUC from 0.507 to 0.827 and AUPR from 0.440 to 0.790. The improvement was smaller on BioSNAP, while performance decreased slightly on BindingDB-AIBind.

We then evaluated compound ranking separately for 101 DUD-E^48^ targets and 14 LIT-PCBA^49^ targets (Fig. 6d,e). Using one reference ligand per target, similarity-band ranking increased the median enrichment factor at the top 1% of ranked compounds (EF 1%) from 0.22 to 23.75 on DUD-E and from 0.15 to 2.27 on LIT-PCBA, relative to unadapted AI-Bind. Library adaptation alone did not improve median EF 1% on either benchmark, despite its improvements in pair prediction. Pocket-based DrugCLIP achieved the highest median EF 1% on both benchmarks, at 31.17 and 3.40, respectively. Performance varied substantially across individual targets.

### Consensus virtual screening prioritizes compounds targeting PD-associated hub proteins

We ranked the pdRGs and Open Targets genes separately using within-module connectivity, increased expression in PD, and ChEMBL pharmacological evidence. The top 20 genes from each list yielded 36 distinct genes. From this shortlist, we selected seven PD-associated proteins for screening: *LRRK2*, *TNK2*, *FYN*, *DYRK1A*, *USP7*, *SNCA*, and *PKM* (Fig. 7a). These targets included kinases and proteins involved in protein homeostasis and metabolism, with different levels of structural and pharmacological evidence.

**Fig. 7.**
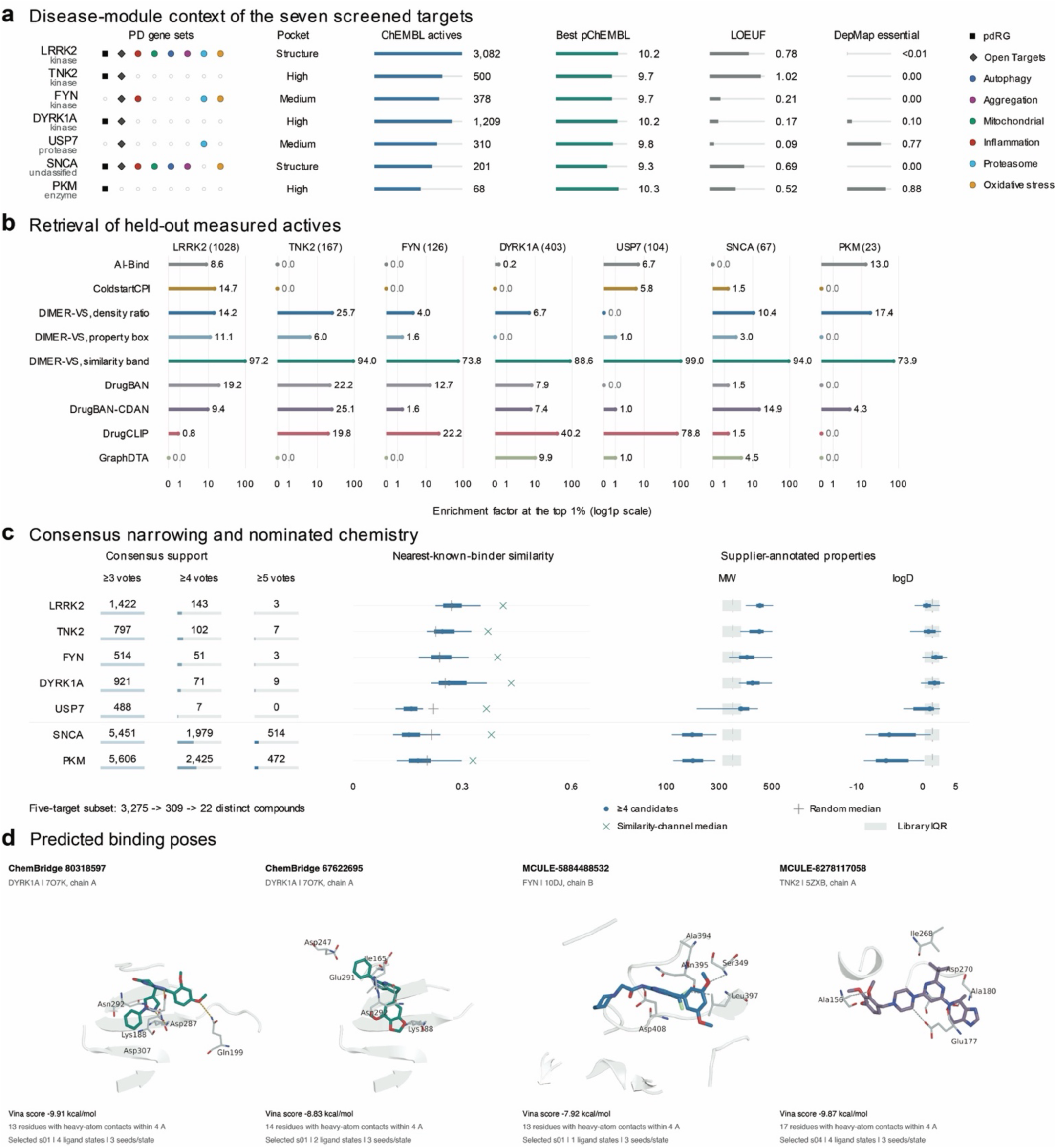
Target-directed virtual screening and consensus candidate nomination. **a,** Disease-module membership and biological, structural, and pharmacological characteristics of screened targets. **b,** Enrichment of held-out known actives when screening 1.82 million compounds against each target. **c,** Candidate counts at different consensus thresholds, with chemical summaries for candidates supported by at least four rankings. Nine ranking configurations each contributed their top 10,000 compounds per target. **d,** Predicted docking poses of four selected candidates in DYRK1A, FYN, and TNK2.

We screened 1,820,535 compounds from ChemBridge^50^ and mcule^51^ using DIMER and five existing prediction models (AI-Bind^14^, ColdstartCPI^15^, DrugBAN^16^, DrugCLIP^17^, and GraphDTA^29^). Performance was evaluated using the same target-specific held-out binders, with separate reference and calibration subsets for similarity-band ranking. Similarity-band ranking achieved the highest EF 1% across all seven targets, ranging from 73.8 to 99.0 (Fig. 7b). Among the other approaches, DrugCLIP performed best on *FYN*, *DYRK1A*, and *USP7*. DIMER-VS density-ratio performed best on *TNK2* and *PKM*. DrugBAN worked best on *LRRK2*, and i*t*s CDAN configuration on *SNCA*.

Since different methods and configurations performed best on different targets (Fig. 7b), we next seek a consensus ranking to prioritize compounds supported by multiple methods. Each configuration contributed one vote for every compound in its top 10,000 for a given target. For the five-target subset comprising *LRRK2*, *TNK2*, *FYN*, *DYRK1A*, and *USP7*, requiring at least three, four, or five votes reduced the candidate set from 3,275 to 309 to 22 distinct compounds (Fig. 7c). Among candidates supported by at least four configurations, those nominated for the five-target subset generally had higher molecular weights and logD values than those nominated for SNCA and PKM. Across all seven targets, consensus candidates had lower median similarity to known binders than candidates prioritized by similarity-band ranking (Fig. 7c).

We used molecular docking to examine the predicted binding poses of four representative candidates from the 22-compound shortlist (Fig. 7d). ChemBridge 80318597 and ChemBridge 67622695 were docked to DYRK1A, MCULE-5884488532 to FYN, and MCULE-8278117058 to TNK2. Their selected poses occupied the ligand-defined pockets, with Vina scores of -9.91, -8.83, -7.92, and -9.87 kcal/mol, respectively.

### Validating the prioritized drug candidates using human midbrain organoid models

We evaluated the top four prioritized drug candidates (polydatin, fedratinib, sorafenib, venlafaxine) from network proximity estimation for their neuroprotective potential by testing in human induced pluripotent stem cells (iPSCs) derived midbrain organoids exposed to 6-hydroxydopamine (6-OHDA) (Fig. 8a; see Methods).

**Fig. 8.**
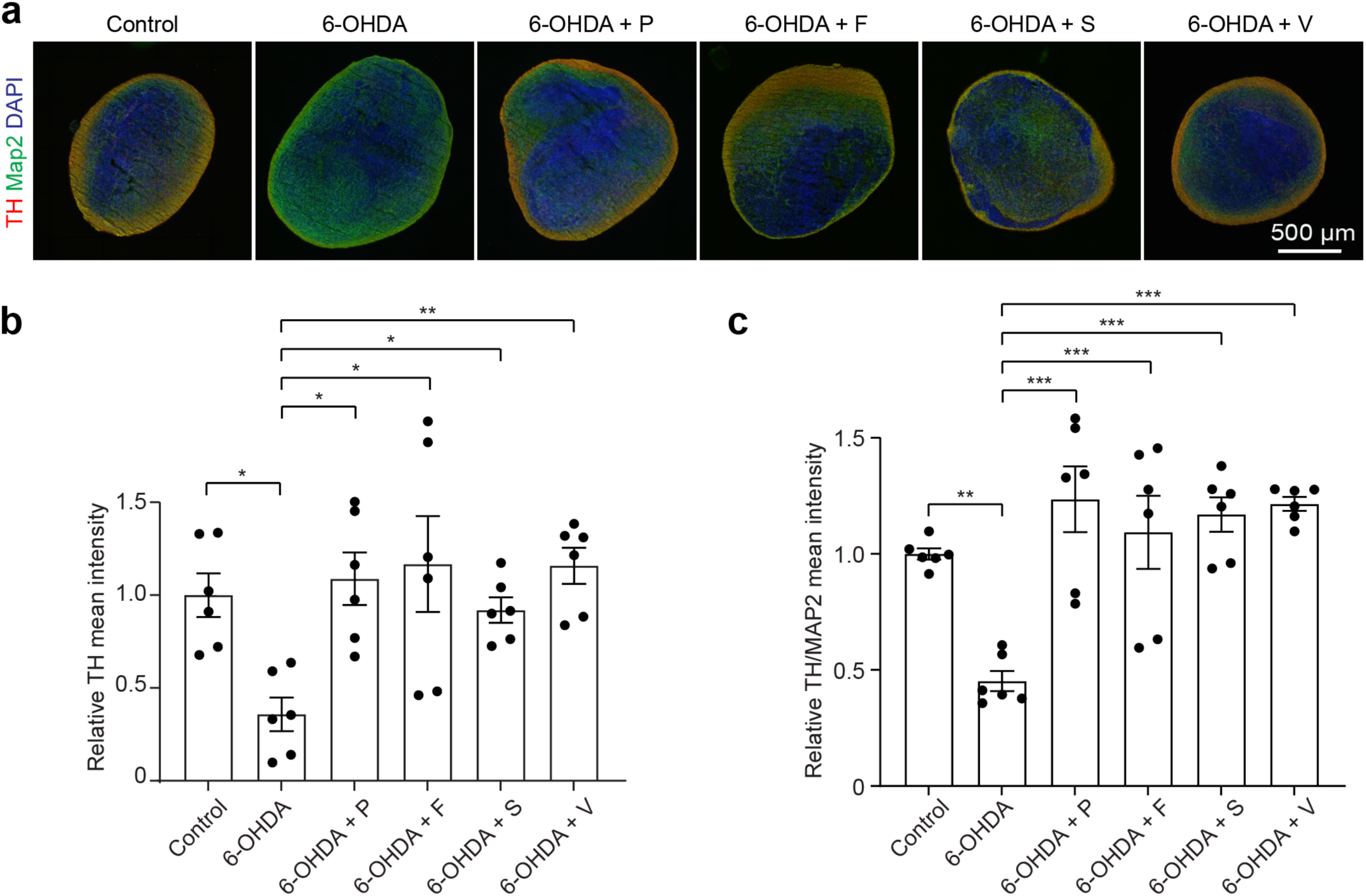
Dopamine neuron protection after compound treatment in midbrain organoids. **a,** Representative immunofluorescence images of human iPSCs-derived midbrain organoids under untreated control conditions, after 6-hydroxydopamine (6-OHDA) exposure, or with 6-OHDA plus polydatin (P; 25 µM), fedratinib (F; 1 µM), sorafenib (S; 1.25 µM), or venlafaxine (V; 1 µM). Compound treatment began one day before 100 µM 6-OHDA exposure, continued during the two-day exposure and for seven subsequent days. Tyrosine hydroxylase (TH), red; MAP2, green; DAPI, blue. Scale bar, 500 µm. **b and c,** Relative TH mean fluorescence intensity (b) and TH/MAP2 mean intensity ratio (c). n = 6 images from 3 organoids per group. * p < 0.05, **p < 0.01, ***p < 0.001.

We labelled dopamine neurons (TH, red), pan-neurons (Map2, green), and nuclei (DAPI, blue) on organoid sections and compared the peripheral TH and TH/Map2 mean intensity (see Methods) as the readouts of the protective ability of the four prioritized drug candidates (Fig. 8b,c). Exposure to 6-OHDA significantly reduced the peripheral TH and TH/MAP2 mean intensity compared to the untreated control, suggesting specific damage to dopamine neurons (Fig. 8b,c). Treatment with each of the four compounds rescued the dopamine neuron injury (Fig. 8b,c). Together, these observations provide experimental support for DIMER’s ability to prioritize compounds with protective potential of dopamine neurons in the human midbrain organoid parkinsonian model.

## Discussion

DIMER connects disease-associated genes to two complementary routes for therapeutic discovery: repurposing compounds with annotated targets and screening new compounds against selected disease-associated proteins. Its contribution lies in using a common biological framework to connect these routes while incorporating information relevant to different stages of candidate selection. In PD, our analyses show that network proximity, transcriptional direction, cellular context, and target-specific screening performance provide distinct information that can change which compounds warrant experimental investigation (Figs. 1-7). The organoid experiments provide initial support for selected repurposing candidates (Fig. 8), while the virtual-screening results nominate additional compounds for biochemical and functional evaluation (Figs. 6-7).

An interesting finding is that existing pharmacology reaches PD-associated genes largely through their interacting proteins. Half of the 36 genes shared across the genetic and mechanism-associated sets lacked direct drug-target annotations in our compendium, and 71% of qualifying drug-module associations involved targets outside the corresponding disease gene set (Fig. 2). Restricting discovery to compounds that directly bind disease-associated proteins would therefore exclude many of the connections identified here. The absence of canonical symptomatic-treatment targets from the eight gene sets further illustrates the distinction between the genetic and mechanistic evidence represented in these sets and the neurotransmitter systems targeted by established therapies.

The transcriptional analyses demonstrate why the direction of a drug’s effect requires separate evaluation. Both established PD treatments and drugs associated with parkinsonism were proximal to PD gene sets, showing that network association can accompany different pharmacological outcomes (Fig. 3). RevERT adds information about whether drug-induced expression changes oppose or reinforce a disease-associated signature. However, this direction depended on the disease reference: whole SNc and dopamine neuron profiles were weakly correlated, and several module genes changed in opposite directions. These differences may reflect changes in tissue composition, responses in non-neuronal populations, and the molecular state of surviving dopamine neurons^44,45,52^. Concordance across references can strengthen prioritization, whereas discordance identifies biological context that requires further investigation. Disease-associated expression changes may also include compensatory responses^53^, so transcriptional reversal should be interpreted as supporting evidence rather than proof of therapeutic benefit.

Cell type-resolved networks further changed the interpretation of drug-disease associations (Fig. 5). Only a subset of static candidates remained proximal in dopamine neurons, while other associations emerged in neuronal and non-neuronal populations. These results suggest that proximity measured in a reference interactome cannot be assumed to apply uniformly across the SNc. Associations involving CSF1R inhibitors in microglia^54^ and S1P receptor modulators in endothelial cells illustrate how cellular context can suggest a site of action^55^. Broadly proximal kinase inhibitors also connected to disease genes through different targets across cell classes. Such information can guide experimental design by identifying which populations and molecular responses should be examined for a given compound. The current networks remain expression-informed models, and direct measurements of protein abundance, activity, and interactions would be needed to improve the proposed cellular mechanisms.

The combination analysis extends this framework to drugs that approach a disease module through different target neighborhoods (Fig. 4). This provides a rationale for investigating whether a candidate adds effects that are complementary to an existing treatment^7^. The association between network separation and documented adverse drug interactions requires careful interpretation: the difference between separated and overlapping pairs was substantially reduced after accounting for shared targets. Target-set separation therefore supports mechanistic prioritization but does not establish combination safety or pharmacological synergy. Likewise, the absence of a listed interaction is not evidence that a combination is safe. The nominated pairs provide a starting point for experiments that measure joint efficacy, dose dependence, and toxicity.

The virtual-screening benchmarks reveal an important distinction between predicting compound-protein pairs and retrieving active compounds early in a ranked library^20^ (Fig. 6). Adapting AI-Bind to the screening library improved pair-classification performance but did not improve median enrichment at the top 1% on the two screening benchmarks. Ligand-informed ranking produced substantially greater improvements in early retrieval, indicating that target-specific chemical evidence can be particularly valuable when only a small number of compounds can be tested. They also define the limits of the present benchmarks: training overlap was permitted in the pair-classification evaluations, so these improvements do not establish generalization to entirely unseen compounds or proteins^56^.

We anticipate that DIMER will be useful for studies seeking to translate genetic discoveries into pharmacological hypotheses. For PD, it provides a way to examine new uses of existing drugs, design complementary additions to established treatments, and prioritize compounds against disease-associated proteins. The underlying components can be adapted to other complex diseases by supplying appropriate gene sets, expression references, and screening libraries. As these resources expand, DIMER can guide focused experimental screens that link molecular mechanisms to candidate interventions.

## Methods

### Overview of the DIMER platform

DIMER (Direct and Indirect Module Evaluation for Repurposing and Screening) integrates network-based drug repurposing with target-directed virtual screening. We applied it to PD using genetic and mechanism-associated gene sets. (1) The network component places disease-associated genes on a human PPI network and evaluates drugs through four analyses: static proximity on the unweighted interactome; RevERT transcriptional reversal; target-set separation for complementary combinations; and cell-type- and condition-resolved proximity using single-nucleus expression data. (2) The virtual-screening component builds upon AI-Bind VecNet, with library-adapted fine-tuning and reference-ligand similarity-band ranking. Complementary sequence- and pocket-based models contribute to benchmark comparisons and consensus candidate nomination.

Network analyses were implemented in R and virtual screening in Python using PyTorch, RDKit and gensim, with an R interface through reticulate. Four selected repurposing compounds were evaluated in human midbrain organoids.

### Parkinson’s disease risk-gene module

The disease gene sets comprised 175 PD risk genes (pdRGs) from Dou *et al.*^4^ and 508 genes supported by genetic evidence in Open Targets^39^. The Open Targets set was further annotated into six overlapping modules using Gene Ontology, Reactome and curated assignments: inflammation, mitochondrial dysfunction, autophagy, protein aggregation, proteasome dysfunction and oxidative stress. Gene identifiers were reconciled through HGNC and mapped to the interactome node labels.

Module coherence on the interactome was assessed by the size of the largest connected component (LCC) of each gene-set-induced subgraph, compared against an empirical null of degree-matched random gene sets of the same size. For each of 1,000 randomizations we drew a size- and degree-matched gene set (matching on binned node degree; see below) and recorded its LCC. We report the observed LCC, the null mean, a standardized effect size *z* = (LCC_obs_ − *μ*_null_)/*σ*_null_, and a one-sided empirical *P* calculated as described under *Statistical analysis*.

### Human protein-protein interaction (PPI) network

Drug-module distances were computed on a literature- and database-derived human interactome represented as an undirected, unweighted graph, comprising 18,223 nodes and 524,156 edges^6^. Self-loops and duplicate edges were removed; static proximity computations were restricted to the largest connected component of 18,208 nodes so that shortest-path lengths are finite between analyzed nodes. Node degree, used for degree-matched null models, was computed on this graph.

### Human drug-target interaction (DTI) network

DTIs were harmonized across seven resources: BindingDB^34^, ChEMBL v34^57^, DrugBank^33^, DrugCentral^35^, the IUPHAR/BPS Guide to Pharmacology^31^, PharmGKB^30^, and the Therapeutic Target Database^36^, into a single non-redundant table. Compounds were unified on DrugBank^33^ identifiers with cross-references to ChEMBL^57^, PubChem^32^, ChEBI^58^, and structure identifiers (InChIKey^59^, InChI^59^, SMILES^60^); protein targets were unified on UniProt^61^ accessions and mapped to the interactome through HGNC identifiers^62^.

Each interaction was annotated into three catalogs: mechanistic, affinity-only, or pharmacogenomic-only. All mapped interactions entered the primary proximity analysis, which included 2,651 drugs with at least one interactome target. Small-molecule status, structure availability, and withdrawal status were recorded per compound. Candidate reporting subsequently applied eligibility and route-evidence checks, including observed distances recalculated after removing pharmacogenomic-only edges, without recomputing the permutation null.

### Annotation of repurposing drugs

We defined 32 drug classes based on mechanism of action, therapeutic use, and drug type, using annotations from ChEMBL v34^57^ and DrugBank^33^. Each drug received one primary class, with manual review to resolve conflicting annotations and maintain consistency across related drugs. Drugs that did not fit any of the 32 categories were assigned to “Other,” giving 33 classes in total. There were 17 classes with fewer than 10 eligible proximal candidates. For data visualization in Fig. 2a, we thus grouped those 17 classes as “17 classes (<10 candidates each).”

### Adverse drug-drug interaction (DDI) reference

To prevent nomination of unsafe combinations, we curated a clinically documented adverse DDIs reference from Cheng et al.^7^ and DDInter 2.0^37^. Combinations were annotated into three catalogs: listed, not listed, or not assessable. These annotations were reported separately from network complementarity; an unlisted pair was not interpreted as established to be safe.

### Single nucleus (sn) reference and differential expression

Cell type- and condition-resolved analyses used SNpc single nucleus RNA-seq data from Kamath et al.^5^ (GSE178265). Nuclei were assigned to the eight major cell classes defined in the original study. Pseudobulk profiles were generated by summing UMI counts within donor, cell type and condition; groups containing fewer than 16 nuclei were excluded. For cell types with at least three donors per condition, edgeR negative-binomial quasi-likelihood models compared PD with controls, adjusting for available donor covariates (age, sex and post-mortem interval).

### Static network proximity

Static proximity used the closest-distance formulation on the unweighted interactome. For a drug *i* with interactome-mapped target set *T_i_* and the module *P*, proximity is the mean shortest-path distance from each target to its nearest module gene,

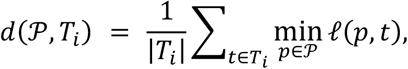

where *l* (⋅,⋅) is the unit-weight shortest-path length.

Statistical significance was assessed against a degree-matched permutation null that randomizes both the module and the drug’s targets.

Nodes were binned into 20 quantile bins of log_10_(degree + 1); each target was replaced by a randomly drawn gene from its own degree bin (without replacement within a bin).

For each permutation we drew a matched random module and matched random target set and recomputed the proximity, yielding a null distribution with mean *μ_i_* and standard deviation *σ_i_*. We report a standardized effect size and a one-sided empirical *P*,

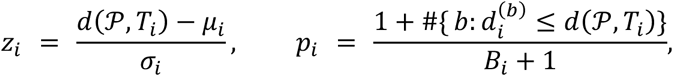

with a negative *z_i_* indicating targets closer to the module than degree-matched expectation and *B_i_* the number of evaluable null draws. We used *B* = 1,000 permutations; drugs with degenerate nulls or non-finite standardized scores were excluded from nomination. Candidate prioritization required *z_i_* ≤ -1.96 and *P* < 0.05.

### RevERT transcriptional reversal score

PD transcriptional directions were estimated separately from laser-captured SNc dopamine neurons^45^ (GSE182622) and whole SNc tissue^44^ (GSE136666). Counts were preprocessed with edgeR and analyzed with limma-voom. The neuronal model adjusted for sex, age and post-mortem interval, with donor correlation estimated using duplicateCorrelation; the tissue model adjusted for sex.

To annotate the predicted direction of a drug’s effect, we adapted the perturbational-directionality concept of Gross *et al.*^6^ using Connectivity Map (CMap)/LINCS L1000 Level-5 compound signatures^43^. DrugBank compounds were mapped to CMap perturbagens using structural identifiers, normalized names and aliases, with mapping provenance retained. We used quality-controlled signatures over measured landmark genes in 11 selected neural or reference cell contexts, retaining all qualifying doses and exposure times. For each disease reference *r*, we defined a PD directional signature from the sign of the case-vs.-control log fold change. For a perturbation signature *k* with CMap values *ζ_kg_*,

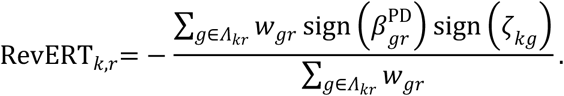

A positive RevERT indicates reversal of the PD-associated direction, and a negative RevERT indicates reinforcement of that direction. Here, *Λ_kr_* contains module genes with evaluable, nonzero disease and perturbation directions; at least three overlapping genes were required. Gene weights *w_gr_* were proportional to -log_10_*P*_g*r*_, rescaled to a maximum of one within each module and reference. Statistical significance used a two-sided normal approximation under independent, zero-mean drug-effect signs, with null standard deviation 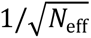 and 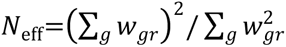. Drug-level scores averaged all qualifying signatures within each disease reference. A consistent direction required at least three signatures, an absolute mean score of at least 0.1 and at least 60% sign agreement.

### Network-based drug-combination

We quantified the network separation between the target sets of two drugs using the framework of Cheng *et al.* Let *T_A_* and *T_B_* denote the sets of distinct interactome-mapped targets of drugs *A* and *B*, respectively, and let *ℓ*(*u*, *v*) denote the shortest-path distance between proteins *u* and *v* in the human interactome. For each drug *X* ∈ {*A*, *B*}, we calculated the mean within-set nearest-neighbor distance:

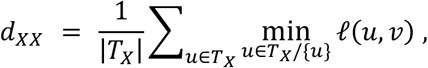

where *d_AA_* and *d_BB_* summarize the distance from each target to its nearest other target of the same drug. Self-distances were excluded from the within-set calculation. For single-target drugs, we assigned a within-set distance of zero by convention.

The symmetric between-set nearest-neighbor distance was calculated as

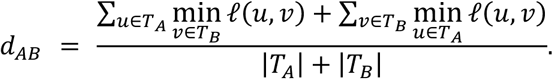

This definition pools the nearest-neighbor distances in both directions, giving each target equal weight. Targets shared by the two drugs contribute zero to the corresponding between-set distances. We then calculated the network separation score:

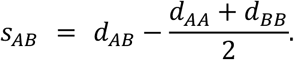

Each unordered pair was assigned to one of six exposure classes from the two proximity scores (*z*<-1.96) and the separation (*s_AB_*≥0, indicating separated target neighborhoods): P1 overlapping exposure, P2 complementary exposure, P3 indirect exposure, P4 single exposure, P5 non-exposure and P6 independent action. P2, in which both drugs reach the module through topologically distinct target neighborhoods, was the prioritized combination class. The stricter nomination subset required positive separation and *P* < 0.05 for both partners, satisfaction of candidate-eligibility and route-evidence checks. Adverse-interaction annotations were retained separately.

### Cell type- and condition-resolved dynamic proximity

The dynamic model incorporates cell-type and disease-condition evidence into the interactome and module. Edges and node anchors are reweighted using the signals below.

#### Context signals

Three normalized signals are computed per gene *g* and cell type *c*. The within-cell-type abundance is the pooled expression fraction,

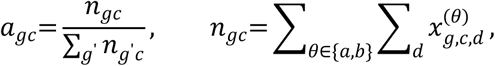

where 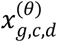 is the pseudobulk count of gene *g* in cell type *c*, donor *d* and condition *θ*, pooled over donors and conditions for stability. The denominator sums over genes in the count matrix. The within-cell-type condition score captures the magnitude of the case-versus-reference perturbation,

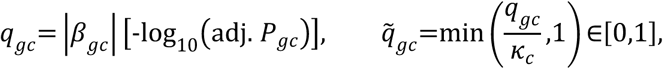

where *κ_c_* is the 99th percentile of the positive raw scores within cell type *c*, capping the influence of extreme genes. Relative cell-type specificity normalizes abundance by the maximum across cell types for a fixed gene,

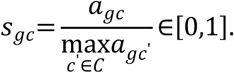

This assigns a value of one to the cell type with the highest abundance of a gene. A signed condition direction 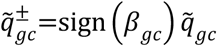 and a condition-specificity partition are retained for interpretation but excluded from the weights.

#### Detection gating

Each signal is gated by the fraction of expressing cells against a threshold *τ*=0.10. The node gate is *γ_gc_*=**1**{*π_gc_*≥*τ*}, where *π_gc_* is the pooled detection fraction; the condition gate requires detection in either state, 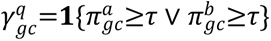. The condition score is set to zero when its gate fails.

#### Node anchor weights

Both module anchors and drug-target anchors receive a gated, convex combination of specificity and condition score,

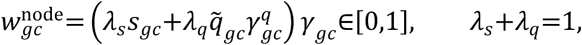

with (*λ_s_*,*λ_q_*)=(0.5, 0.5) for module anchors and (0.7,0.3) for drug targets. The additive form retains an anchor that is strong on either axis; when an axis is inactive its coefficient is removed and the remainder renormalized to sum to one.

#### Dynamic edge layer

An interactome edge is retained in a cell type only if both endpoints pass the pooled detection gate. Retained edges are assigned bounded lengths based on detection fractions,

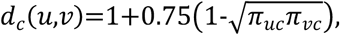

so that consistently detected interactions are shorter. The node set is fixed across cell types; only the open-edge set and edge lengths vary with *c*.

#### Weighted module distance and drug proximity

Let *M* ⊆ *P* be the module anchors with positive weight in cell type *c*. We augment the graph with one auxiliary source s joined to every anchor *g*∈*M* by an edge whose length penalizes low-weight anchors,

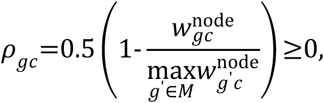

and run a single shortest-path computation. The distance from the weighted module to any node *v* is then, in closed form,

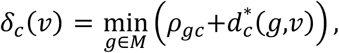

where 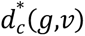 (*g*,*v*) is the weighted shortest-path length in the dynamic graph. For drug *i* with positive-weight targets 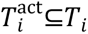 reachable from the module and target weights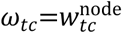, the context proximity is the target-weighted mean module distance,

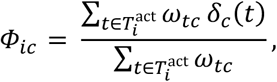

with smaller *Φ_ic_* indicating a drug whose well-expressed targets lie close to the weighted disease module in cell type *c*.

#### Significance

As in the static analysis, significance was assessed by a context-matched permutation null using up to 20 quantile bins per matched covariate. The primary null matched on degree, abundance and cell-type specificity; a sensitivity null additionally matched on the condition score. Matching bins were relaxed progressively when no unused match was available. For each of 1,000 permutations we drew a matched random module, preserving its anchor weights, and a matched random target set, using the weights of the sampled target nodes, then recomputed *δ_c_* and *Φ_ic_*. With null mean *μ_ic_* and standard deviation *σ_ic_*,

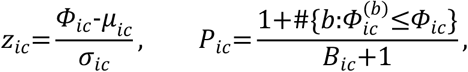

where *B_ic_* is the number of evaluable null draws. Non-finite standardized scores and degenerate nulls were excluded from nomination. Prioritization used the same rule as the static analysis.

### DIMER virtual screening model

DIMER virtual screening builds on the VecNet configuration of AI-Bind^14^ with library adaptation and ligand-similarity integration, as described below.

#### Featurization

A compound was represented from its SMILES string by the set of Morgan substructure identifiers up to radius 2, embedded and summed into a 300-dimensional Mol2vec vector, with an unknown-token fallback for unseen substructures,

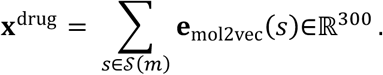

A protein target was represented from its amino-acid sequence by non-overlapping 3-grams embedded and summed into a 100-dimensional ProtVec vector,

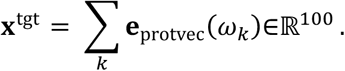

Both raw vectors were standardized by frozen training-set constants (mean and scale) that define the coordinate system of the pretrained weights throughout adaptation and inference.

#### Architecture and scoring

Each modality is projected to a 2,048-dimensional hidden representation and concatenated, and a multilayer perceptron head returns a binding logit:

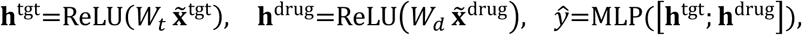

with the head mapping 4096→512→512→1 through ReLU nonlinearities. The binding score is the mean sigmoid output over a five-fold model ensemble,

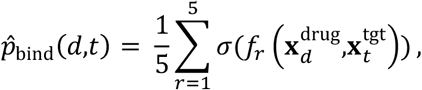

where *σ* is the logistic function and *f_r_* the *r*-th fold. Structure featurization is cached, so a full library is featurized once and scored against any number of targets.

#### Library-adapted fine-tuning

To specialize the baseline to a user’s chemical space, we optionally fine-tuned each fold on labeled drug-target pairs restricted to the library’s chemical subspace. In the property-box scheme, the subspace is the union of the library’s InChIKey set and its physicochemical envelope, i.e. an axis-aligned box on molecular weight, topological polar surface area, hydrogen-bond donors and acceptors, and calculated logP, and labeled pairs from the training splits falling inside it are retained.

The density-ratio scheme weighted labeled training compounds to match the library’s chemical distribution. A class-balanced logistic classifier distinguished library compounds from unique training compounds using 12 molecular descriptors and up to 20,000 sampled library compounds. For estimated library-membership probability *q*(*x*), the raw weight was *q*(*x*)/(1-*q*(*x*)). Within each fold, weights were divided by their median, clipped to 0.1-10 and rescaled to mean one. Both schemes used library chemical features without library binding labels. Fine-tuning minimized weighted binary cross-entropy,

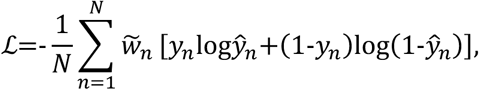

with *w̃_n_*=1 for retained pairs in the property-box scheme and the conditioned importance weight in the density-ratio scheme. Optimization used AdamW (learning rate 10^-4^, weight decay 10^-2^, batch size 256), while freezing the target-projection tower and normalization constants and updating only the drug-projection tower and head. A stratified 20% validation split selected the lowest weighted validation loss, including the pretrained state, with patience two and at most 20 epochs.

#### Similarity-band ranking

For targets with known ligands, the maximum Tanimoto similarity *s_it_* between compound *i* and the reference set was calculated using 2,048-bit Morgan fingerprints of radius 2. Let *u_it_* be the percentile of the sequence-model score within the screened set, scaled to [0, 1]. For band width 0<*b*<1, the combined score was

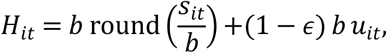

where the second term is scaled by (1 − *ε*), so that it is strictly smaller than one band width and the sequence percentile can only order compounds within a band. We set *ε* = 10^−3^ during our study.

Similarity determines the order between bands, and the model score orders compounds within a band. At *b*=0, ranking uses similarity alone; at *b*=1, it uses the sequence-model percentile alone. Where calibration data were available, *b* was selected from 0, 0.005, 0.01, 0.02, 0.05, 0.10, 0.20, 0.50 and 1 to maximize enrichment in the top 1%, resolving ties toward the larger band. The public default retrieves binders by exact protein-sequence matching to a ChEMBL36^57^ reference bundle and retains density-adapted sequence ranking when reference or calibration support is insufficient.

### Benchmark virtual screening models

Compound-protein classification on BindingDB^34^, the AI-Bind BindingDB collection^14^, and BioSNAP^47^ used full deduplicated pair sets and reported the area under the receiver operating characteristic curve (ROC AUC) and average precision. The evaluation used the existing model weights on these pair collections; training overlap was permitted. Similarity-assisted comparisons used common scored pairs for reference-covered proteins, with both the full reference sets and references excluding canonical structures appearing in the corresponding evaluation pairs. Screening evaluation used the matched set of 101 DUD-E^48^ and 14 LIT-PCBA^48^ targets, reporting ROC AUC, BEDROC (*α*=80.5) and enrichment factors at 0.5%, 1% and 5%.

Comparators were AI-Bind^14^, ColdstartCPI^15^, DrugBAN^16^, DrugBAN-CDAN^16^, DrugCLIP^17^ and GraphDTA^29^. Adaptation used unlabeled screening compounds; the similarity-band benchmark used the pretrained sequence model, one crystallographic reference ligand and fixed *b*=0.01.

### Hub-gene prioritization for virtual screening

Hub genes were ranked separately within the 175-pdRG and 508-gene Open Targets sets. For each set, we constructed its induced subgraph on the undirected PPI network and calculated module degree, defined as the number of interacting genes in the same set. All genes were retained, including zero-degree genes. The hub percentile was 100 times the average rank of module degree divided by the number of genes in the set, rounded to one decimal place. Betweenness centrality, k-core membership, articulation status, LCC membership and global degree were retained as descriptive annotations.

The final score equaled the hub percentile plus ten points for each of three binary evidence flags: increased expression in PD, binding evidence and association with a clinical-stage compound. Increased expression required positive PD-versus-control change at adj. *P* < 0.05 in at least one of three bulk references (DA-neuron RNA-seq, whole-SN RNA-seq, or SN microarray meta-analysis) or eight SNpc cell-class pseudobulk contrasts. Binding evidence required at least one ChEMBL IC50, Ki or Kd record at pChEMBL ≥ 6. ChEMBL targets were resolved through UniProt accessions. The clinical-stage flag required a compound with maximum clinical phase ≥ 1 among the retrieved target-activity records (pChEMBL ≥ 5), or a clinical-stage compound with a curated mechanism involving the target.

Genes were ordered by descending final score, then module degree. The top 20 genes from each list were combined into a 36-gene shortlist. Seven targets were selected from this shortlist following review of biological context, measured ligand evidence and available ligand-bound structures. Pocket annotations, loss-of-function constraint, DepMap essentiality and clinical-trial annotations provided additional context for this selection. The selected targets were LRRK2, TNK2, FYN, DYRK1A, USP7, SNCA and PKM.

### Virtual-screening application to PD-associated genes

LRRK2, TNK2, FYN, DYRK1A, USP7, SNCA and PKM were screened against 1,820,535 ChemBridge^50^ and mcule^51^ compounds. Human single-protein ChEMBL binders with IC50, Ki or Kd measurements at pChEMBL ≥ 6 were split into reference, calibration and held-out thirds, using at most 500 reference binders. Similarity bands were calibrated against the unlabeled library background. Methods were evaluated on the same held-out binders, with compound coverage recorded. For production screening, an internal half split selected the band before references were drawn from the full binder pool.

Consensus nomination used eight production-ranking runs: AI-Bind^14^, ColdstartCPI^15^, DIMER-VS density ratio, DIMER-VS property box, DrugBAN^16^, DrugBAN-CDAN^16^, DrugCLIP^17^ and GraphDTA^29^. Each contributed its top 10,000 compounds per target, and candidates were summarized at thresholds of at least three, four and five votes. Similarity-band rankings were analyzed separately. Compound identities were reconciled before voting. Model-family support grouped AI-Bind with the two DIMER adaptations and DrugBAN with DrugBAN-CDAN; the remaining methods each formed a separate family. Chemical summaries used maximum Tanimoto similarity to known binders and supplier-annotated molecular properties, with random library samples as references.

### Docking of consensus compound candidates

The four representative consensus candidates were docked to ligand-defined sites in DYRK1A (PDB 7O7K, chain A), FYN (10DJ, chain B) and TNK2 (5ZXB, chain A). Receptor preparation retained the selected protein chain, removed waters and other nonprotein components, repaired missing side-chain atoms with PDBFixer and added hydrogens and Gasteiger charges with Meeko 0.7.1. Observed heavy-atom coordinates were preserved; unresolved loops were left unmodeled. Candidate preparation enumerated unspecified carbon stereocenters and Molscrub 0.3.2 protonation/tautomer states at pH 7.4. Each state received 20 RDKit ETKDGv3 conformers; the lowest-energy converged MMFF94s conformer was used for docking.

AutoDock Vina 1.2.5 used a rigid receptor, exhaustiveness 32 and three seeds (20261025, 20261026 and 20261027), retaining up to 20 poses per run. Boxes were centered on the crystallographic ligand bounds, with 6 Å padding per face and a minimum dimension of 24 Å. The displayed pose was the lowest-scoring pose across states and seeds, with ties resolved by state identifier, seed and pose rank. Native ligands were redocked from de novo conformers; symmetry-aware heavy-atom root-mean-square deviation (RMSD) was measured without fitting to the native pose. Top-ranked control RMSDs were 1.40-1.43 Å for DYRK1A, 1.57-1.58 Å for FYN and 2.41-3.07 Å for TNK2. The TNK2 structure contained an unresolved 272-292 segment adjacent to the pocket.

Heavy-atom contacts used a 4-Å cutoff. Hydrogen bonds required donor-acceptor distance ≤ 3.5 Å, hydrogen-acceptor distance ≤ 2.5 Å and donor-hydrogen-acceptor angle ≥ 120°, using RDKit donor/acceptor assignments. Selected poses were rendered in PyMOL 3.1.0.

### Human midbrain organoid culture and drug screening

#### Culture and maintenance of human midbrain organoids

Human midbrain organoids were cultured from human induced pluripotent stem cells (hiPSCs, Gibco Cat# A19845) using the STEMdiff™ Midbrain Organoid Differentiation Kit (STEMCELL Technologies, Cat# 100-1096) according to the manufacturer’s protocol. hiPSCs were maintained in mTeSR plus medium (STEMCELL Technologies, Cat# 100-1130) and were routinely tested for mycoplasma contamination. We used hiPSC cells with normal pluripotent stem cell morphology and less than 5% differentiation for organoid culture. We generated cell aggregates in AggreWell™800 (STEMCELL Technologies, Cat# 34811), then cultured them in the formation, expansion, and differentiation media supplied with the STEMdiff™ Midbrain Organoid Differentiation Kit. Beginning on day 43, organoids were maintained in STEMdiff™ Neural Organoid Maintenance Medium (STEMCELL Technologies; Cat# 100-0120). Organoids were maintained under static conditions at 37°C in a humidified incubator containing 5% CO₂.

#### Drug treatment

We used 100 µM 6-hydroxydopamine hydrobromide (6-OHDA, Sigma-Aldrich, Cat. No. 162957) treatment for two days, followed by additional culture for 7 days to induce dopamine neuron damage. To assess the compound effects on potential rescue of dopamine neurons, we first pretreat the organoids at day 56 with polydatin (MCE, Cat. No. HY-N0120A, 25 µM), fedratinib (MCE, Cat. No. HY-10409, 1 µM), sorafenib (MCE, Cat. No. HY-10201, 1.25 µM), and venlafaxine (MCE, Cat. No. HY-B0196, 1 µM) for one day. In the following two days, we treated the organoids with 100µM 6-OHDA together with the corresponding compounds. We then maintained the organoids for another 7 days with the compounds only.

#### Immunofluorescence staining

On day 56, organoids were fixed with 4% paraformaldehyde (Poly Scientific R&D Crop; SKU S2334) at 4°C overnight, cryoprotected in 30% sucrose solution, embedded in 7.5% gelatin solution, and sectioned into 20 µm sections with a Leica CM1900 Cryostat. Organoid sections were incubated with primary antibodies against tyrosine hydroxylase (TH, rabbit, Sigma-Aldrich, Cat. Ab152) and microtubule-associated protein 2 (MAP2, guinea pig, SYSY, Cat.188004), followed by incubation with species-specific secondary antibodies conjugated to Alexa Fluor 647 (donkey anti-rabbit, Invitrogen, Cat. A32795) and Alexa Fluor 488 (Goat anti-guinea pig, Fisher Scientific, Cat. 188004), respectively. Nuclei were counterstained with DAPI (Sigma-Aldrich, Cat. D9542-5MG). Sections were mounted with Prolong mounting medium (Thermo Fisher Scientific, Cat. P36930) and were stored at 10°C before imaging.

#### Imaging and quantification

Fluorescence images were acquired using Echo Revolution Fluorescence Microscope with a 20x objective lens. Imaging parameters were calibrated using the positive control group and the identical parameters were maintained across the groups. Fluorescence signal intensity was quantified using Fiji (ImageJ). For each organoid section, a region of interest (ROI) was manually drawn along the outer boundary of the organoid. An inner ROI was then generated by shrinking the original ROI by 70 µm using the Enlarge function. The annular region between the outer and inner ROIs was obtained using the XOR operation. Background fluorescence was measured from adjacent region without detectable fluorescence signal and subtracted from the mean fluorescence intensity of the annular ROI to obtain the background-corrected mean fluorescence intensity. Both TH and MAP2 fluorescence intensities were quantified for each organoid section from the same annular ROI.

### Statistical analysis

Permutation-based analyses used 1,000 resamples; empirical *P* was calculated as the number of evaluable null statistics at least as extreme as observed plus one, divided by the number of evaluable null draws plus one. Tests were one-sided toward shorter proximity or larger LCCs. Standardized effect sizes (*z*) were reported alongside empirical *P*. Multiple testing was controlled by the Benjamini-Hochberg procedure within each differential-expression contrast and each network module, separately for each dynamic null model, with adjusted results denoted adj. *P*. Candidate prioritization used the stated nominal thresholds; adjusted results were reported separately. RevERT used the analytic null described above; exploratory comparisons of imputed RevERT used two-sided Mann-Whitney tests.

## Data and code availability

Code and curated reference data are available at https://github.com/JHELab2024/network_based_drug_discovery_for_PD.

